# Localization of S1P_1_ Receptor Signaling in the rat, mouse and human Central Nervous System

**DOI:** 10.1101/2024.01.29.577735

**Authors:** Jonatan Martínez-Gardeazabal, Gorka Pereira-Castelo, Marta Moreno-Rodríguez, Alberto Llorente-Ovejero, Manuel Fernández, Iván Fernández-Vega, Iván Manuel, Rafael Rodríguez-Puertas

**Affiliations:** Department of Pharmacology, Faculty of Medicine and Nursing. University of the Basque Country (UPV/EHU), B° Sarriena s/n, 48940 Leioa, Spain; Neurodegenerative Diseases, BioCruces Bizkaia Health Research Institute, 48903 Barakaldo, Spain; Department of Neurology, Hospital Universitario de Cruces, 48903 Barakaldo, Spain; Department of Pathology, Hospital Universitario Central de Asturias, Avda. Roma, s/n, 33011 Oviedo, Spain; Health Research Institute of Principality of Asturias (ISPA), Av. del Hospital Universitario, s/n, 33011 Oviedo, Spain

**Keywords:** S1P_1_ receptor, human, rodents, brain, [^35^S]GTPγS, mapping

## Abstract

Some specific lipid molecules present in the brain are signaling molecules at the intracellular compartments or behaving as neurotransmitters or neuromodulators of other systems, binding to specific G protein-coupled receptors (GPCR) for neurolipids. One of these receptors is the sphingosine 1-phosphate receptor subtype 1, coupled to G_αi/o_-proteins and involved in cell proliferation, growth or neuroprotection. Thus, an interesting target for neurodegenerative diseases, such as Alzheimer’s. The present study compares the human cerebral distribution of the activity mediated by S1P_1_ receptor with the that in the brain of rodent experimental models, rat and mice by functional autoradiography, measuring the [^35^S]GTPγS binding stimulated by the S1P_1_ receptor selective agonist CYM-5442 to get the anatomy of the S1P_1_ receptor activity.

The S1P_1_ receptor-mediated activity is, together with that of the CB_1_ cannabinoid receptor, one of the highest recorded for any GPCR in most of the grey matter areas of the brain, reaching up to 50%-500% over basal, depending on the agonist and brain area. The S1P_1_ receptor signaling is very relevant in those areas that regulate learning and memory processes, such as the basal forebrain, but also in others involved in control of motor processes or nociception e.g. basal ganglia. The results also reveal that the rat would be a preferable experimental model to extrapolate for the S1P_1_ receptor-mediated responses in human brain.

## 1. Introduction

Some of the endogenous lipid-based signaling molecules are considered as neurotransmitters with agonist-like or neuromodulatory properties, which can be denominated neurolipids (similar to the term “neuropeptides”). Neurolipids are a particular class of neurotransmitters since they are originated from membrane lipid precursors by the action of different enzymes such as phospholipases or sphingomyelinases, among others (reviewed in Manuel et al., 2020). Neurolipids are synthesized on demand in a calcium-dependent manner from membrane lipid precursors and not stored in presynaptic vesicles, and can be metabolized both enzymatically and non-enzymatically by oxidative degradation (Shimizu, 2009). The distribution or activity of neurolipid receptors for the endocannabinoid system has been described (Herkenham et al., 1991), and also for the lysophosphatidic acid (LPA) (González De San Román et al., 2015) or the sphingosine 1-phosphate (S1P) (Jiang et al., 2021). S1P was firstly described as a lysophospholipid acting through endothelial differentiation gene (EDG) GPCRs family. Later this family of receptors was differentiated in S1P and LPA receptors, thus, EDG-1 was renamed as S1P_1_ (Chun et al., 2002; Hla et al., 2001; Lee et al., 1998). The expression of the *s1pr1* gene has been observed in many tissues, being especially abundant in spleen, brain, heart, lung, liver and thymus (Zhang et al., 1999). Intracellularly, the S1P_1_ receptor transduces its signaling through the activation of G_αi/o_ proteins, which leads to a wide variety of biological responses, such as effects on the immune response, the regulation of the cellular barrier integrity, cell proliferation, migration, proliferation, and angiogenesis (Ben Shoham et al., 2012; Camp et al., 2020; Garcia et al., 2001; Liu et al., 2019; Matloubian et al., 2004; Pyne and Pyne, 2017; Spiegel and Weinstein, 2004). In this regard, the activity of the S1P_1_ receptor has been studied by the [^35^S]GTPγS assay in rat tissue using S1P and in mouse measuring the stimulation mediated by the partial agonist for S1P_1_, SEW2871 (Sim-Selley et al., 2018, 2009; Waeber and Chiu, 1999), and the density of the S1P_1_ receptor has recently been described using a new radioligand assay with [^3^H]CS1P1 (Jiang et al., 2021). The pharmacological study of S1P receptors in the CNS made a leap with the development of fingolimod, a high affinity agonist for S1P_1_, S1P_3_, S1P_4_ and S1P_5_ receptors after its *in vivo* phosphorylation, which is able to down-regulate S1P_1_, so this compound is also considered a “functional S1P_1_ antagonist” (Brinkmann et al., 2002; Gräler and Goetzl, 2004).

Thus, the present study is aimed to analyze the anatomical brain distribution of functional coupling of G_αi/o_ proteins to S1P_1_ receptors in samples of representative brain areas enriched in S1P_1_ receptors: frontal cortex, temporal cortex, caudate-putamen, hippocampus, amygdala, nucleus basalis of Meynert and cerebellum of *postmortem* human brain. The comparison of the human brain S1P_1_ activity with that in rat and mice models has been also studied to validate these species for models of neurological diseases such as multiple sclerosis and Alzheimer’s type dementia.

## 2. Materials and Methods

### 2.1. Subjects and animals

#### 2.1.1. *Postmortem* human brain samples

*Postmortem* human brain samples were obtained from different cerebral tissue banks: Biobank of the Basque Country and tissue bank of the Asturias Central University Hospital. The samples were obtained at autopsy after getting informed consent in accordance with the ethics committees of the University of the Basque Country (UPV/EHU) (CEISH/244MR/2015/RODRIGUEZ PUERTAS), following the Code of Ethics of the World Medical Association (Declaration of Helsinki), and warranting the privacy rights of the human subjects. The samples (n=8) had not shown any evidence of metabolic or neurological disease, and after the neuropathological study, no abnormality was observed in the brain. The brain samples were immediately frozen at −80°C after autopsy. The mean of age and *postmortem* time are shown in the following table (Table 1).

**Table 1.** Demographic data from human samples. Data are mean ± SEM.

| n |  |  | Age<br>(years) | *Postmortem*<br>(hours) |
| --- | --- | --- | --- | --- |
| Total | Male | Female |  |  |
| 8 | 5 | 3 | 74 ± 4.7 | 9 ± 2.3 |

#### 2.1.2. Sprague-Dawley rats

Two months old male Sprague-Dawley rats (200–250 g) were used for the study (n=8). The animals were housed four per cage (50 cm length x 25 cm width x 15 cm height) at a temperature of 22°C and in a humidity-controlled (65%) room with a 12:12 hours light/dark cycle, with access to food and water *ad libitum.* Every effort was made to minimize animal suffering and to use the minimum number of animals. All procedures were carried out in accordance with European animal research laws (Directive 2010/63/EU) and the Spanish National protocols that were approved by the Local Ethical Committee for Animal Research of the University of the Basque Country (CEEA 388/2014, CEEA M20-2018-52/54).

#### 2.1.3. Swiss mice

Two month old male Swiss mice (25–35 g) (n=8), were also used for histochemical and autoradiographic studies. The animals were housed four per cage (29 cm length x 16 cm width x 12 cm height) at a temperature of 22°C and in a humidity-controlled (65%) room with a 12:12 hours light/dark cycle, with access to food and water *ad libitum.* Local Ethical Committee for Animal Research of the University of the Basque Country (CEEA 366-1/2014, CEEA M20/2018/192).

#### 2.1.4. Animal fresh tissue

All animals (Swiss mice and Sprague-Dawley rats) used in the present study were anesthetized with ketamine/xylazine (90/10 mg/kg; IP) and then euthanized by decapitation. The brain samples were quickly removed by dissection (4°C), fresh frozen and kept at −80°C. In order to process the samples, fresh frozen tissues were raised to −20°C, and 20 µm thick slices were cut in a cryostat (Microm HM550, Walldorf, Germany) and mounted onto gelatin-coated slides. These slices were kept at −25°C until used in autoradiographic studies or histochemical methods.

The present study covers a wide number of areas of the rat and mouse brain. So, the most representative brain levels, both in coronal and sagittal slices, were used to measure the activity of the S1P_1_ receptor activity in each of the areas under study according to the areas described in the rat and mouse stereotaxic coordinates atlases (Paxinos and Franklin, 2001; Paxinos and Watson, 2013).

### 2.2. Labeling of activated G_αi/o_ proteins by [^35^S]GTP**γ**S binding assay

Preliminary studies were performed to determine the experimental conditions for detection of S1P_1_ receptor activity by [^35^S]GTPγS assay. S1P is a lipid ligand, thus we followed a similar protocol to those previously used for detection of activity mediated by other lipid neurotransmitters, such as CB_1_ receptors (Manuel et al., 2014). Regarding the ligand used for the stimulation, we utilized the S1P endogenous ligand, which possess affinity for all described subtypes of S1P receptors; and the agonist CYM-5442, that has a higher affinity for S1P_1_ receptors according to the IUPHAR/BPS. The advantage of using CYM-5442 over S1P lies in that it allows us to evaluate uniquely the activity of S1P_1_ receptor subtype. Moreover, in order to test the specificity of CYM-5442 for the S1P_1_ receptor, the specific S1P_1_ receptor antagonist W146 antagonized the stimulation exerted by CYM-5442. All these assays were performed in sagittal rat brain slices (Figures S1 and S2).

Briefly, fresh 20 μm sections obtained from rodent brain samples and *postmortem* human brain samples were air dried, followed by two consecutive incubations in HEPES-based buffer (HEPES 50 mM, NaCl 100 mM, MgCl_2_ 3 mM, EGTA 0.2 mM and BSA 0.5%; pH 7.4) for 30 min at 30°C to remove the endogenous ligands. Slices were incubated for 2h at 30°C in the same buffer but supplemented with GDP 2 mM, DTT 1 mM and 0.04 nM [^35^S]GTPγS. Basal binding was determined in two consecutive slices in the absence of the agonist. For the agonist-stimulated binding measurement, another consecutive slice was incubated with the same reaction buffer, yet in the presence of CYM-5442. The specific S1P_1_ receptor antagonist W146 (10 μM) together with CYM-5442 (10 μM) was used in consecutive slices to ascertain that the activation was mediated by that specific receptor subtype. Non-specific binding was defined by competition with GTPγS (10 μM) in another consecutive section. Then, slices were washed twice in cold (4°C) 50 mM HEPES buffer (pH 7.4), dried and exposed to β-radiation sensitive film with a set of [^14^C]-standards calibrated for ^35^S. After 48h the films were developed, scanned and quantified by transforming optical densities into nCi/g tissue equivalent (nCi/g t.e.) units using a calibration curve defined by the known values of the [^14^C]-standards (FIJI Software, Bethesda, MD, USA). Background and non-specific binding were subtracted from all experimental conditions. Then, the net stimulations were calculated by subtracting the basal binding.

### 2.4. Histochemical methods: Thionine staining

Thionine staining was performed to facilitate the identification of neuroanatomical structures. Adjacent sections to those used in [^35^S]GTPyS autoradiography were stained with thionine. Tissues were mounted onto gelatin-coated slides and hydrated after thawing. The hydration was performed by immersing tissues for 5 minutes in ethanol solutions in descending order (100%, 96%, 70% and 50%). Then sections were submerged in thionine solution for 5 minutes. Later, tissues were washed with deionized water, and introduced in ethanol solutions (50%, 70%, 96% and 100%) to dehydrate the tissue and covered with DPX as the mounting medium.

## 3. Results

### 3.1. Localization of S1P_1_ receptor-mediated activity by [^35^S]GTPγS autoradiography assay in the rat CNS

First, the S1P_1_ receptor-mediated activity was measured by [^35^S]GTPγS autoradiography assay in rat brain. For this purpose, brain slices were collected rostro-caudally every 500 µm to obtain an exhaustive mapping of the S1P_1_ receptor stimulation distribution in the different nucleus and areas present in the rat brain. The obtained net stimulations of [^35^S]GTPγS binding by CYM-5442 (10 μM), obtained by subtracting basal stimulation, showed that the activity of S1P_1_ receptor was widely distributed and abundant along all the brain, showing specific patterns corresponding to the anatomical distribution of active S1P_1_ receptors in rat brain (Figure 1).

**Figure 1.**
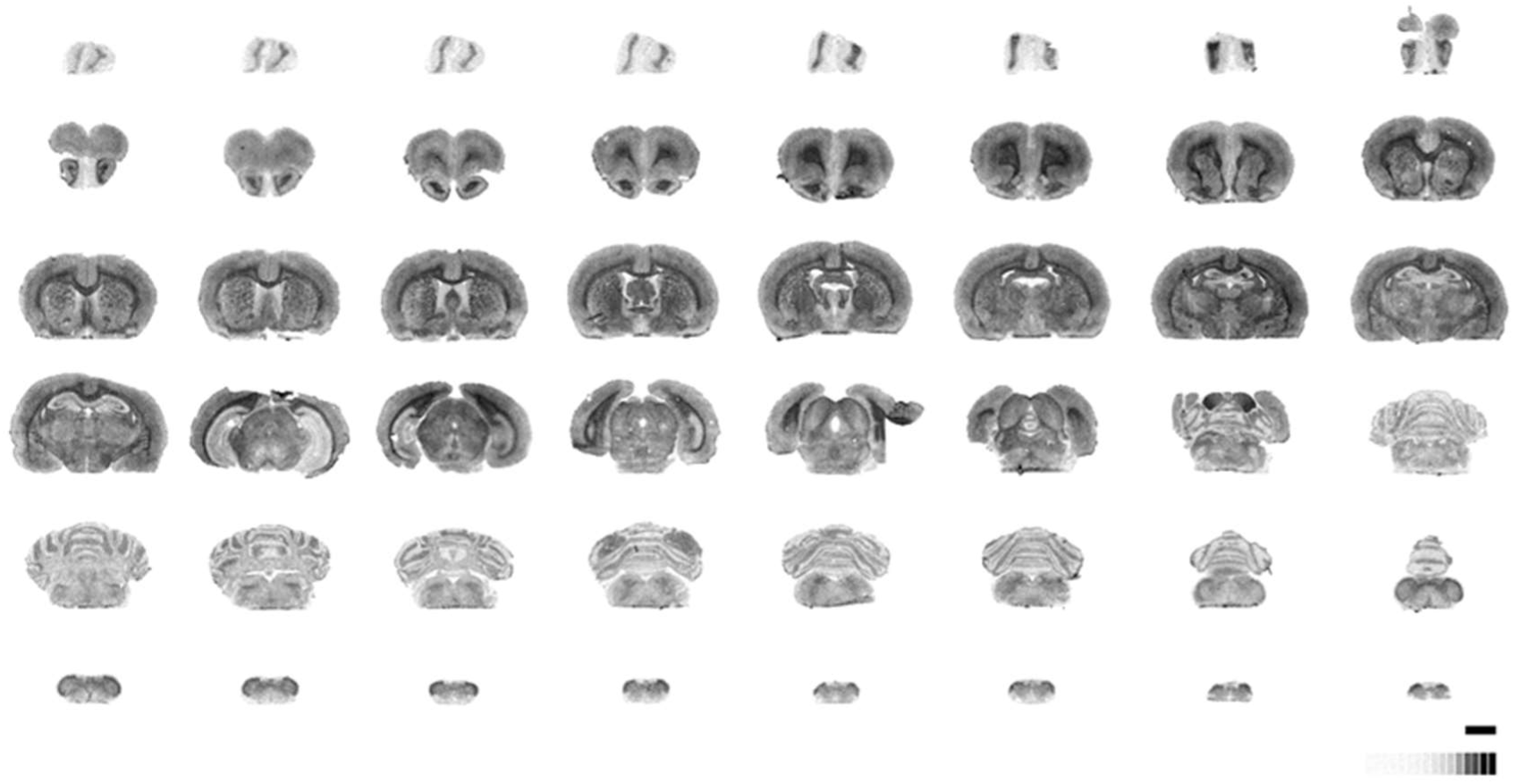
Representative autoradiograms of [^35^S]GTPγS binding following the stimulation with the specific agonist of S1P_1_ receptor, CYM-5442 (10 μM), in whole rat brain. Each of the brain slices was collected rostro-caudally every 500 µm. [^14^C]-Standard (0-35000 nCi/g t.e.). Scale bar = 6 mm.

The activity mediated by S1P_1_ receptors was quantified in 139 different brain areas throughout the entire rat brain, from the olfactory bulb to the spinal cord. The measurement of the S1P_1_ activity revealed the basal nucleus magnocellularis as the brain nucleus where the stimulation was one of the highest (2889 ± 194 nCi/g t.e.), while the middle cerebellar peduncle was the area with the lowest stimulation (171 ± 173 nCi/g t.e.) (Figure 2). Taking the stimulation in these structures as a reference, the following ranges of receptor stimulation (nCi/g t.e.) intensities were stablished: very abundant, >2300; abundant, 2300-1700; moderate, 1700-1100; sparse, 1100-580; and very sparse, <580.

**Figure 2.**
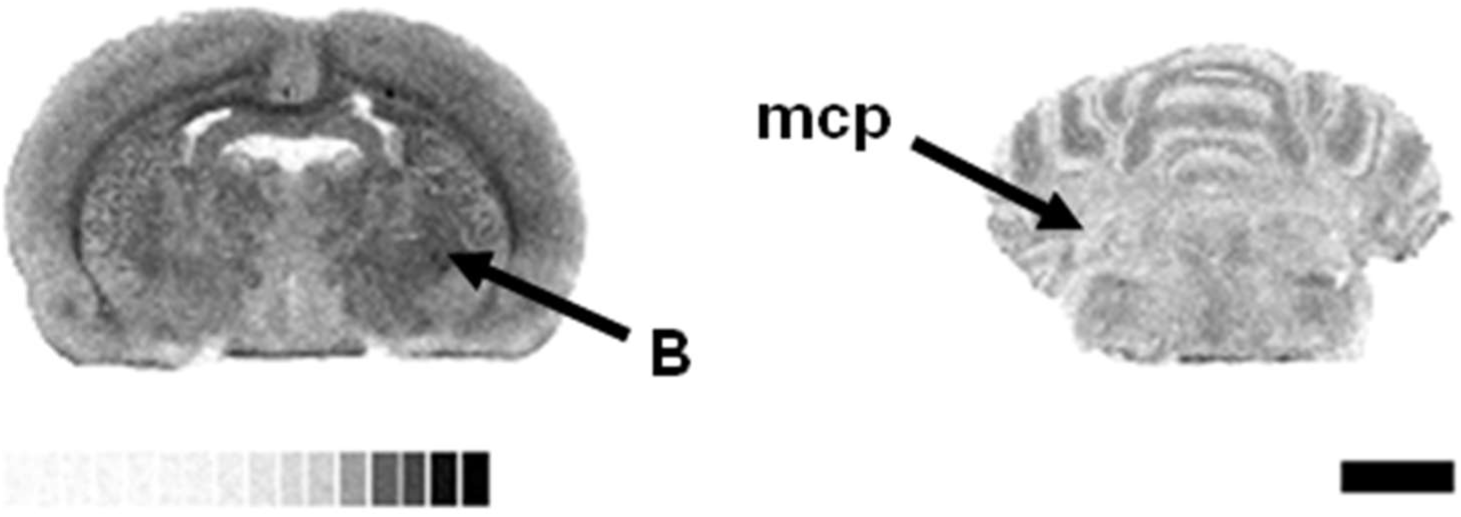
Detail of representative autoradiograms of [^35^S]GTPγS stimulated by the specific agonist of S1P_1_ receptor, CYM-5442 (10 μM), in rat brain showing the high binding at nucleus basalis magnocellularis (B), and the low activity at the middle cerebellar peduncle (mcp). [^14^C]-Standard (0 – 35000 nCi/g t.e.) Scale bar = 6 mm.

In the olfactory areas, one of the highest stimulations mediated by S1P_1_ receptor was found in the rostral migratory stream (1682 ± 162 nCi/g t.e.). The lateral olfactory part and the anterior olfactory area showed a sparse S1P_1_ receptor activity (1078 ± 104 nCi/g t.e. and 1006 ± 105 nCi/g t.e., respectively). On the contrary, both the plexiform, internal and external, and glomerular layers had one of the lowest stimulations in this area (Table 2).

**Table 2.**
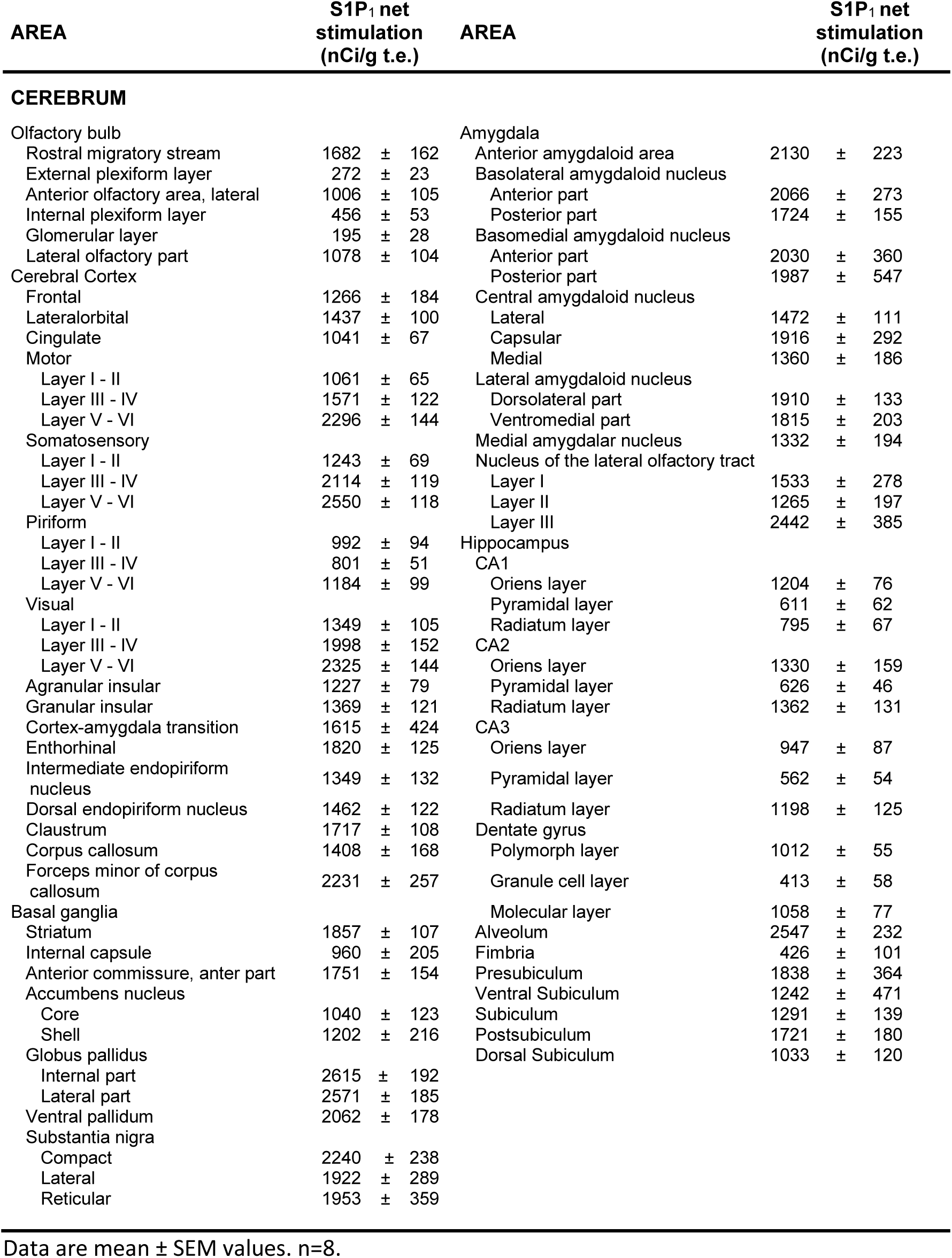
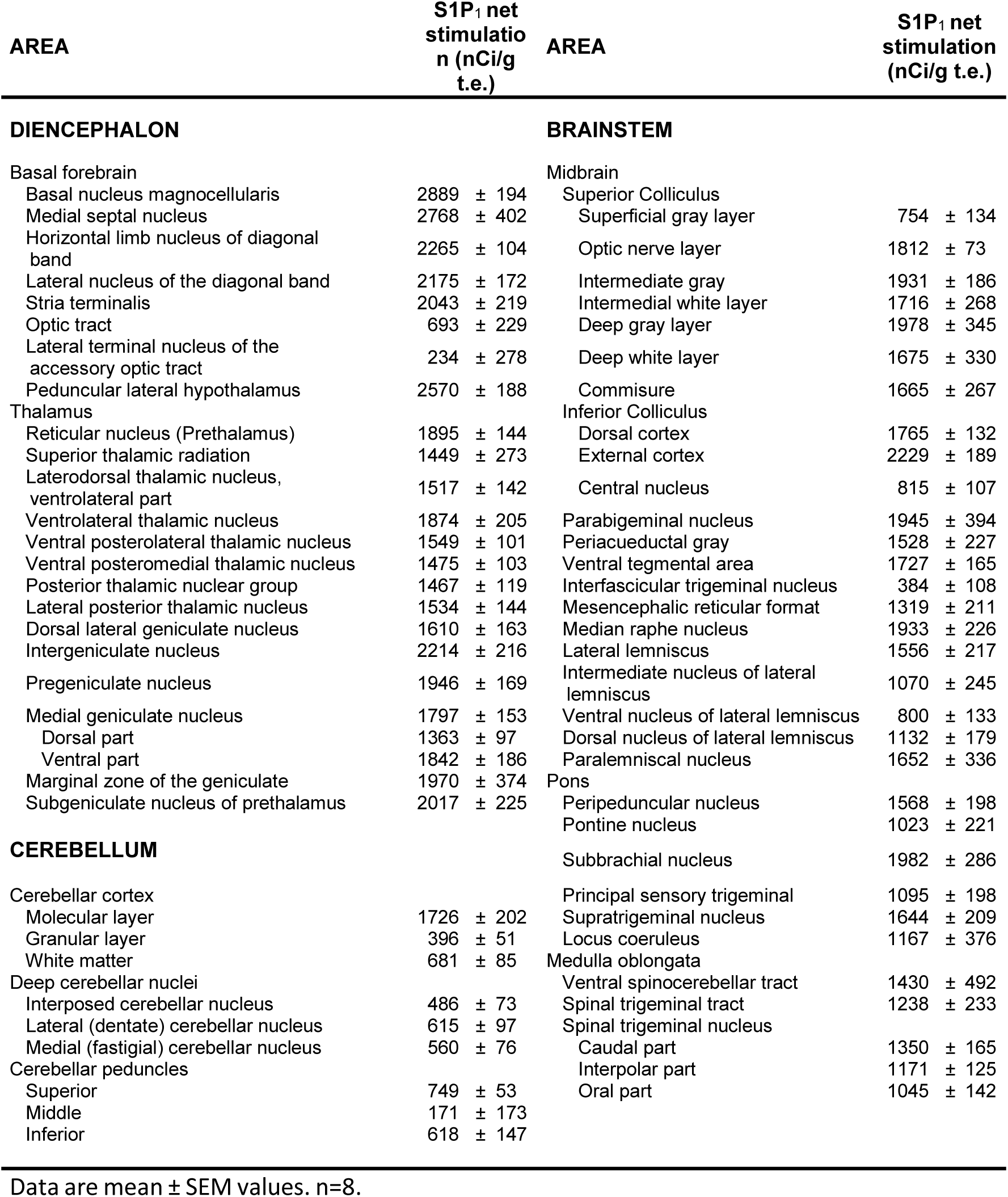
Net stimulation of [^35^S]GTPγS binding by the agonist of S1P_1_ receptor, CYM-5442 (10 μM), in rat brain (nCi/g t.e).

| AREA | S1P<sub>1</sub> net stimulation (nCi/g t.e.) | AREA | S1P<sub>1</sub> net stimulation (nCi/g t.e.) |
| --- | --- | --- | --- |
| **CEREBRUM** |  |  |  |
| Olfactory bulb |  | Amygdala |  |
| Rostral migratory stream | 1682 ± 162 | Anterior amygdaloid area | 2130 ± 223 |
| External plexiform layer | 272 ± 23 | Basolateral amygdaloid nucleus |  |
| Anterior olfactory area, lateral | 1006 ± 105 | Anterior part | 2066 ± 273 |
| Internal plexiform layer | 456 ± 53 | Posterior part | 1724 ± 155 |
| Glomerular layer | 195 ± 28 | Basomedial amygdaloid nucleus |  |
| Lateral olfactory part | 1078 ± 104 | Anterior part | 2030 ± 360 |
| Cerebral Cortex |  | Posterior part | 1987 ± 547 |
| Frontal | 1266 ± 184 | Central amygdaloid nucleus |  |
| Lateral orbital | 1437 ± 100 | Lateral | 1472 ± 111 |
| Cingulate | 1041 ± 67 | Capsular | 1916 ± 292 |
| Motor |  | Medial | 1360 ± 186 |
| Layer I - II | 1061 ± 65 | Lateral amygdaloid nucleus |  |
| Layer III - IV | 1571 ± 122 | Dorsolateral part | 1910 ± 133 |
| Layer V - VI | 2296 ± 144 | Ventromedial part | 1815 ± 203 |
| Somatosensory |  | Medial amygdalar nucleus | 1332 ± 194 |
| Layer I - II | 1243 ± 69 | Nucleus of the lateral olfactory tract |  |
| Layer III - IV | 2114 ± 119 | Layer I | 1533 ± 278 |
| Layer V - VI | 2550 ± 118 | Layer II | 1265 ± 197 |
| Piriform |  | Layer III | 2442 ± 385 |
| Layer I - II | 992 ± 94 | Hippocampus |  |
| Layer III - IV | 801 ± 51 | CA1 |  |
| Layer V - VI | 1184 ± 99 | Oriens layer | 1204 ± 76 |
| Visual |  | Pyramidal layer | 611 ± 62 |
| Layer I - II | 1349 ± 105 | Radiatum layer | 795 ± 67 |
| Layer III - IV | 1998 ± 152 | CA2 |  |
| Layer V - VI | 2325 ± 144 | Oriens layer | 1330 ± 159 |
| Agranular insular | 1227 ± 79 | Pyramidal layer | 626 ± 46 |
| Granular insular | 1369 ± 121 | Radiatum layer | 1362 ± 131 |
| Cortex-amygdala transition | 1615 ± 424 | CA3 |  |
| Entorhinal | 1820 ± 125 | Oriens layer | 947 ± 87 |
| Intermediate endopiriform nucleus | 1349 ± 132 | Pyramidal layer | 562 ± 54 |
| Dorsal endopiriform nucleus | 1462 ± 122 | Radiatum layer | 1198 ± 125 |
| Clastrum | 1717 ± 108 | Dentate gyrus |  |
| Corpus callosum | 1408 ± 168 | Polymorph layer | 1012 ± 55 |
| Forceps minor of corpus callosum | 2231 ± 257 | Granule cell layer | 413 ± 58 |
| Basal ganglia |  | Molecular layer | 1058 ± 77 |
| Striatum | 1857 ± 107 | Alveolum | 2547 ± 232 |
| Internal capsule | 960 ± 205 | Fimbria | 426 ± 101 |
| Anterior commissure, anter part | 1751 ± 154 | Presubiculum | 1838 ± 364 |
| Accumbens nucleus |  | Ventral Subiculum | 1242 ± 471 |
| Core | 1040 ± 123 | Subiculum | 1291 ± 139 |
| Shell | 1202 ± 216 | Postsubiculum | 1721 ± 180 |
| Globus pallidus |  | Dorsal Subiculum | 1033 ± 120 |
| Internal part | 2615 ± 192 |  |  |
| Lateral part | 2571 ± 185 |  |  |
| Ventral pallidum | 2062 ± 178 |  |  |
| Substantia nigra |  |  |  |
| Compact | 2240 ± 238 |  |  |
| Lateral | 1922 ± 289 |  |  |
| Reticular | 1953 ± 359 |  |  |

Data are mean ± SEM values. n=8.

| AREA | S1P<sub>1</sub> net stimulation (nCi/g t.e.) | AREA | S1P<sub>1</sub> net stimulation (nCi/g t.e.) |
| --- | --- | --- | --- |
| **DIENCEPHALON** |  | **BRAINSTEM** |  |
| Basal forebrain |  | Midbrain |  |
| Basal nucleus magnocellularis | 2889 ± 194 | Superior Colliculus |  |
| Medial septal nucleus | 2768 ± 402 | Superficial gray layer | 754 ± 134 |
| Horizontal limb nucleus of diagonal band | 2265 ± 104 | Optic nerve layer | 1812 ± 73 |
| Lateral nucleus of the diagonal band | 2175 ± 172 | Intermediate gray | 1931 ± 186 |
| Stria terminalis | 2043 ± 219 | Intermedial white layer | 1716 ± 268 |
| Optic tract | 693 ± 229 | Deep gray layer | 1978 ± 345 |
| Lateral terminal nucleus of the accessory optic tract | 234 ± 278 | Deep white layer | 1675 ± 330 |
| Peduncular lateral hypothalamus | 2570 ± 188 | Commisure | 1665 ± 267 |
| Thalamus |  | Inferior Colliculus |  |
| Reticular nucleus (Prethalamus) | 1895 ± 144 | Dorsal cortex | 1765 ± 132 |
| Superior thalamic radiation | 1449 ± 273 | External cortex | 2229 ± 189 |
| Laterodorsal thalamic nucleus, ventrolateral part | 1517 ± 142 | Central nucleus | 815 ± 107 |
| Ventrolateral thalamic nucleus | 1874 ± 205 | Parabigeminal nucleus | 1945 ± 394 |
| Ventral posterolateral thalamic nucleus | 1549 ± 101 | Periacqueductal gray | 1528 ± 227 |
| Ventral posteromedial thalamic nucleus | 1475 ± 103 | Ventral tegmental area | 1727 ± 165 |
| Posterior thalamic nuclear group | 1467 ± 119 | Interfascicular trigeminal nucleus | 384 ± 108 |
| Lateral posterior thalamic nucleus | 1534 ± 144 | Mesencephalic reticular format | 1319 ± 211 |
| Dorsal lateral geniculate nucleus | 1610 ± 163 | Median raphe nucleus | 1933 ± 226 |
| Intergeniculate nucleus | 2214 ± 216 | Lateral lemniscus | 1556 ± 217 |
| Pregeniculate nucleus | 1946 ± 169 | Intermediate nucleus of lateral lemniscus | 1070 ± 245 |
| Medial geniculate nucleus | 1797 ± 153 | Ventral nucleus of lateral lemniscus | 800 ± 133 |
| Dorsal part | 1363 ± 97 | Dorsal nucleus of lateral lemniscus | 1132 ± 179 |
| Ventral part | 1842 ± 186 | Paralemniscal nucleus | 1652 ± 336 |
| Marginal zone of the geniculate | 1970 ± 374 | Pons |  |
| Subgeniculate nucleus of prethalamus | 2017 ± 225 | Peripeduncular nucleus | 1568 ± 198 |
|  |  | Pontine nucleus | 1023 ± 221 |
| **CEREBELLUM** |  | Subbrachial nucleus | 1982 ± 286 |
| Cerebellar cortex |  | Principal sensory trigeminal | 1095 ± 198 |
| Molecular layer | 1726 ± 202 | Supratrigeminal nucleus | 1644 ± 209 |
| Granular layer | 396 ± 51 | Locus coeruleus | 1167 ± 376 |
| White matter | 681 ± 85 | Medulla oblongata |  |
| Deep cerebellar nuclei |  | Ventral spinocerebellar tract | 1430 ± 492 |
| Interposed cerebellar nucleus | 486 ± 73 | Spinal trigeminal tract | 1238 ± 233 |
| Lateral (dentate) cerebellar nucleus | 615 ± 97 | Spinal trigeminal nucleus |  |
| Medial (fastigial) cerebellar nucleus | 560 ± 76 | Caudal part | 1350 ± 165 |
| Cerebellar peduncles |  | Interpolar part | 1171 ± 125 |
| Superior | 749 ± 53 | Oral part | 1045 ± 142 |
| Middle | 171 ± 173 |  |  |
| Inferior | 618 ± 147 |  |  |

Data are mean ± SEM values. n=8.

Regarding the cerebral cortex, the stimulations were maintained throughout the different areas of the cortex: somatosensory (1883 ± 104 nCi/g t.e.), visual (1874 ± 125 nCi/g t.e.), entorhinal (1820 ± 125 nCi/g t.e.), cortex-amygdala transition (1615 ± 424 nCi/g t.e.), motor (1465 ± 71 nCi/g t.e.), lateral-orbital (1437 ± 100 nCi/g t.e.), granular-insular (1369 ± 121 nCi/g t.e.), frontal (1266 ± 184 nCi/g t.e.), agranular-insular (1227 ± 79 nCi/g t.e.), cingulate (1041 ± 67 nCi/g t.e.) and piriform (984 ± 62 nCi/g t.e.) cortex. Within the different regions of the cortex, it was observed that the stimulations of layers V and VI (mostly layer VI) were higher than layers III and IV (mostly layer IV) and these, in turn, more than layers I and II. Other analyzed nuclei with moderate levels were the dorsal endopiriform nucleus (1462 ± 122 nCi/g t.e.), intermediate endopiriform nucleus (1349 ± 132 nCi/g t.e.) and claustrum (1717 ± 108 nCi/g t.e.). It should be noted, that in some white matter areas the stimulation of the S1P_1_ receptor was high, e.g. in the corpus callosum (1408 ± 168 nCi/g t.e.) and in the minor forceps of corpus callosum (2231 ± 257 nCi/g t.e.), that apparently were areas with high recordings, however the high basal levels in absence of S1P_1_ agonist account for a considerable amount of that labelling (see figure S2).

In the basal ganglia, we found a high activity in the globus pallidus in both internal and lateral part (2615 ± 192 nCi/g t.e. and 2571 ± 185 nCi/g t.e., respectively). Other close areas, such as the ventral pallidum, showed high stimulations (2062 ± 178 nCi/g t.e.). Another structure with elevated stimulations was the substantia nigra, in which the compact part (2240 ± 238 nCi/g t.e.) showed a higher activity than reticular (1953 ± 359 nCi/g t.e.) or lateral part (1922 ± 289 nCi/g t.e.). A high stimulation was also measured in the striatum (1857 ± 107 nCi/g t.e.), delineating the fibers inside (2533 ± 221 nCi/g t.e.). However, the nucleus *accumbens* core and the internal capsule were more sparsely stimulated (1040 ± 123 nCi/g t.e. and 960 ± 205 nCi/g t.e., respectively).

In the amygdala and surrounding structures, the activity of S1P_1_ receptor was abundant and quite homogenous, but it was notably high on layer III of the nucleus of the lateral olfactory tract (2442 ± 385 nCi/g t.e.). Additionally, the anterior part of the basolateral and basomedial amygdaloid nucleus (2066 ± 273 nCi/g t.e. and 2030 ± 360 nCi/g t.e., respectively) showed higher stimulations than their posterior parts (1724 ± 155 nCi/g t.e. and 1987 ± 547 nCi/g t.e.). In the lateral amygdaloid nucleus, both dorsolateral and ventromedial part, were similarly stimulated (1910 ± 133 nCi/g t.e. and 1815 ± 203 nCi/g t.e.). However, in central amygdaloid nucleus a stronger stimulation was measured in capsular part (1916 ± 292 nCi/g t.e.) than in lateral or medial part (1472 ± 111 nCi/g t.e. and 1360 ± 186 nCi/g t.e, respectively).

In the hippocampus, we found a stimulation pattern that was similar in CA1, CA2 and CA3 regions. In all of these regions, the *radiatum* and *oriens* layers showed higher S1P_1_ activity than the pyramidal layer. In the dentate gyrus, the pattern showed that molecular layer and polymorphic layer were stimulated better than granular layer, where very sparse activity was found. It is noteworthy that the *alveolum* was strongly stimulated (2547 ± 232 nCi/g t.e.).

Concerning the basal forebrain, as mentioned, the basal nucleus magnocellularis showed probably the highest specific S1P_1_ activity that we were able to measure (2889 ± 194 nCi/g t.e.). Septal related structures as medial septal nucleus, horizontal limb nucleus of diagonal band and lateral nucleus of the diagonal band showed also high S1P_1_ activity (2768 ± 402 nCi/g t.e.; 2265 ± 104 nCi/g t.e.; 2175 ± 172 nCi/g t.e., respectively). In contrast, lateral terminal nucleus of the accessory optic tract and optic tract were sparsely stimulated (234 ± 278 nCi/g t.e. and 693 ± 229 nCi/g t.e., respectively).

In the thalamus, the S1P_1_ activity was more homogenous than in other regions, but we were able to differentiate some structures as different nucleus of geniculate nucleus or thalamic nucleus.

In cerebellum, we found abundant levels of S1P_1_ activity in the molecular layer of the cerebellar cortex (1726 ± 202 nCi/g t.e.), while the granular layer of the cerebellar cortex showed low S1P_1_-mediated activity (396 ± 51 nCi/g t.e.), in a similar way to cerebellar white matter (681 ± 85 nCi/g t.e.) and inside deep nuclei: dentate (615 ± 97 nCi/g t.e.), fastigial (560 ± 76 nCi/g t.e.) and interposed (486 ± 73 nCi/g t.e.). The cerebellar peduncles were also measured with few or very few stimulations in the superior (749 ± 53 nCi/g t.e.), middle (171 ± 173 nCi/g t.e.) and inferior (618 ± 147 nCi/g t.e.) peduncles.

The midbrain was moderately stimulated, but we identified some structures that showed a specific distribution. In the colliculus, both inferior and superior, we found a pattern in the distribution of the activation of the S1P_1_ receptor. In the inferior colliculus, the central nucleus was poorly stimulated (815 ± 107 nCi/g t.e.) compared to the external cortex or dorsal cortex (2229 ± 189 nCi/g t.e.; 1765 ± 132 nCi/g t.e.). In the superior colliculus, we were able to differentiate the layers that form the superior colliculus, where the superficial layers showed lower levels than optic nerve layer that was highly activated (1812 ± 73 nCi/g t.e). The intermediate and deep layers were strongly stimulated. About the lateral lemniscus, some nucleus were measured with weak activity as dorsal (1132 ± 179 nCi/g t.e.), intermediate (1070 ± 245 nCi/g t.e.) and ventral (800 ± 133 nCi/g t.e.) nucleus, yet lateral lemniscus and paralemniscal nuclei were moderately stimulated (1556 ± 217 nCi/g t.e.; 1652 ± 336 nCi/g t.e.). Other structures were identified with high activity, such as parabigeminal nucleus (1945 ± 394 nCi/g t.e.), median raphe nucleus (1933 ± 226 nCi/g t.e.) and ventral tegmental area (1727 ± 165 nCi/g t.e.); moderate activity, as the periaqueductal gray (1528 ± 227 nCi/g t.e.) and mesencephalic reticular formation (1319 ± 211 nCi/g t.e.); and low stimulation, as interfascicular trigeminal nucleus (384 ± 108 nCi/g t.e.).

Regarding the pons, we showed that the stimulation in this area was lower than in other areas and was more focused in some nucleus as the subbrachial nucleus (1982 ± 286 nCi/g t.e.), supratrigeminal nucleus (1644 ± 209 nCi/g t.e.) or peripeduncular nucleus (1568 ± 198 nCi/g t.e.). In other nucleus of the brain stem moderate stimulations were found, as in locus coeruleus (1167 ± 376 nCi/g t.e.), principal sensory trigeminal (1095 ± 198 nCi/g t.e.) or pontine nucleus (1023 ± 221 nCi/g t.e.).

Finally, in the medulla oblongata, the activity of S1P_1_ receptor was moderate and focused on the spinal trigeminal tract (1238 ± 233 nCi/g t.e.), ventral spinocerebellar tract (1430 ± 492 nCi/g t.e.) and spinal trigeminal nucleus, which could differentiate in the caudal part (1350 ± 165 nCi/g t.e.), interpolar part (1171 ± 125 nCi/g t.e.) and oral part (1045 ± 142 nCi/g t.e.).

### 3.2 Localization of S1P_1_ receptor-mediated activity by [^35^S]GTPγS autoradiography assay in the mouse CNS

The S1P_1_ receptor-mediated activity was also mapped by [^35^S]GTPγS autoradiography assay in mouse brain. In the same way as it was done with the rat, slices of the brain rostro-caudally cut every 500 µm were collected to obtain an exhaustive mapping of the S1P_1_ receptor stimulation in the different structures present in the mouse brain. The obtained stimulations of [^35^S]GTPγS binding by CYM-5442 (10 μM) showed that S1P_1_ receptor activities were high along the mouse brain but were slightly weaker than those observed in the rat brain (Figure 3). These differences are discussed in section 3.3 of this article. For further information about the activity of S1P_1_ receptor in the mouse brain, see the Supplementary Material section (Table S2).

**Figure 3.**
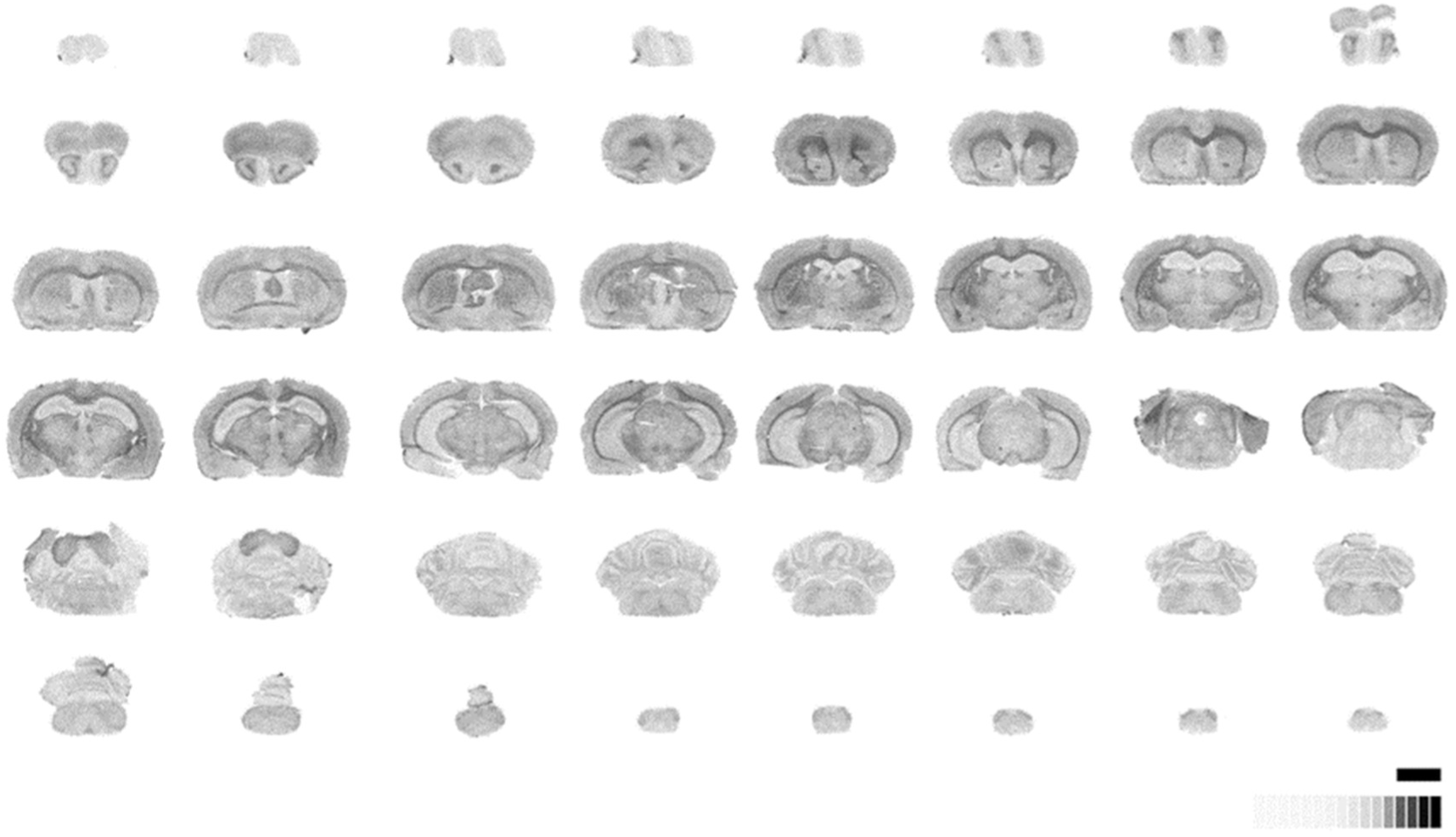
Representative autoradiograms of [^35^S]GTPγS stimulated by specific agonist of S1P1 receptor, CYM-5442 (10 μM) in mouse brain. [^14^C]-Standard (0-35000 nCi/g t.e.). Scale bar = 6 mm.

### 3.3 S1P_1_ receptor activity in human *postmortem* brain

The activity of the S1P_1_ receptor was also analyzed in *postmortem* human brain tissue, exploring the distribution of this receptor in the CNS and comparing it with the most common animal models used in neuropharmacological research, i.e., rat and mouse.

Note the high activity of S1P_1_ receptors in all of the human brain areas that were analyzed. The stimulation pattern of S1P_1_ receptor in human tissue is similar to the activity pattern previously observed in rat brain. In the cortical areas, we observed that the layers III – VI were more stimulated than layers I – II and V – VI in frontal (Layer I – II: 937 ± 154 nCi/g t.e.; Layer III – IV: 1329 ± 254 nCi/g t.e.; Layer V – VI: 1301 ± 175 nCi/g t.e.; figure 4A) and temporal cortex (Layer I – II: 835 ± 141 nCi/g t.e.; Layer III – IV: 920 ± 194 nCi/g t.e.; Layer V – VI: 796 ± 161 nCi/g t.e.; figure 4G), but this pattern changes in entorhinal cortex, where the layers V – VI were highly stimulated compared to layers I – II and III – IV (Layer I – II: 869 ± 160 nCi/g t.e.; Layer III – IV: 863 ± 178 nCi/g t.e.; Layer V – VI: 1394 ± 312 nCi/g t.e.; figure 4F), and in periamygdalar cortex, where all the layers were stimulated homogeneously (Layer I – II: 915 ± 190 nCi/g t.e.; Layer III – IV: 890 ± 234 nCi/g t.e.; Layer V – VI: 886 ± 234 nCi/g t.e.; figure 4D). The white matter areas underneath to the different cortices were weakly stimulated (figures 4E, 4G; Table 3).

**Figure 4.**
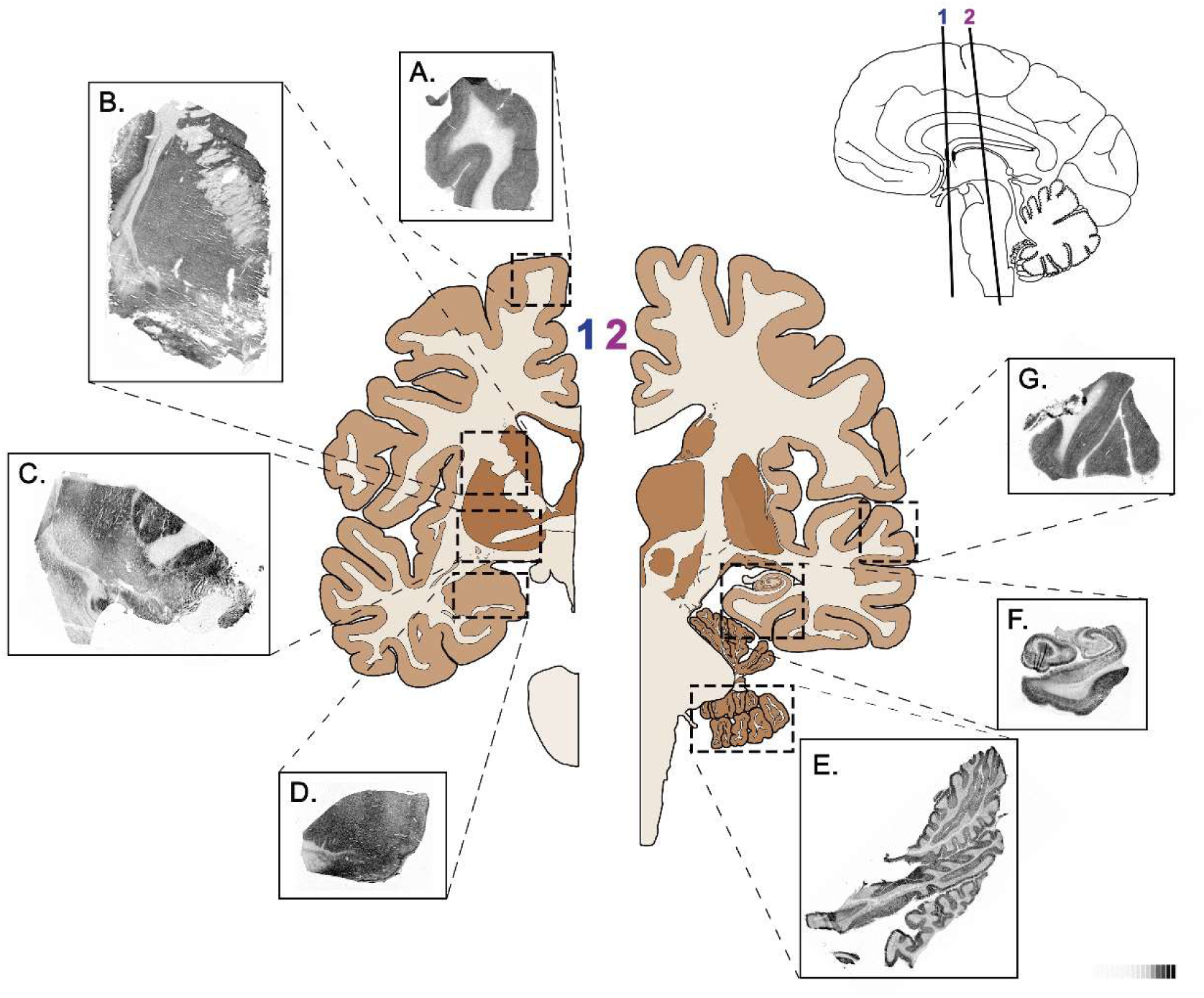
Representative autoradiograms of [^35^S]GTPγS stimulated by specific agonist of S1P_1_ receptor, CYM-5442 (10 μM), in human frontal cortex (A), striatum (B), basal forebrain (C), amygdala (D), cerebellum (E), hippocampus (F) and temporal cortex (G). [^14^C]-Standard (0-35000 nCi/g t.e.).

**Table 3.** Net [^35^S]GTPγS binding stimulated by the specific agonist of S1P_1_ receptor CYM-5442 (10 μM) in human brain (nCi/g t.e.).

| Brain area | Basal<br>(nCi/g t.e.) | S1P<sub>1</sub> net<br>stimulation<br>(nCi/g t.e.) | Brain area | Basal<br>(nCi/g t.e.) | S1P<sub>1</sub> net<br>stimulation<br>(nCi/g t.e.) |
| --- | --- | --- | --- | --- | --- |
| Frontal Cortex |  |  | Hippocampus |  |  |
| Layer I - II | 132 ± 23 | 937 ± 154 | CA1 |  |  |
| Layer III - IV | 156 ± 26 | 1329 ± 254 | Oriens | 165 ± 41 | 1200 ± 291 |
| Layer V - VI | 181 ± 29 | 1301 ± 175 | Pyramidal | 152 ± 41 | 462 ± 91 |
| White Matter | 78 ± 25 | 229 ± 75 | Radiatum | 193 ± 37 | 1438 ± 324 |
| Temporal Cortex |  |  | CA2 |  |  |
| Layer I - II | 277 ± 39 | 835 ± 141 | Oriens | 163 ± 41 | 1210 ± 193 |
| Layer III - IV | 296 ± 43 | 920 ± 194 | Pyramidal | 145 ± 37 | 406 ± 67 |
| Layer V - VI | 339 ± 56 | 796 ± 161 | Radiatum | 189 ± 45 | 1362 ± 275 |
| White Matter | 130 ± 15 | 50 ± 26 | CA3 |  |  |
| Entorhinal Cortex |  |  | Oriens | 205 ± 40 | 1365 ± 264 |
| Layer I - II | 168 ± 49 | 869 ± 160 | Pyramidal | 142 ± 40 | 424 ± 76 |
| Layer III - IV | 185 ± 60 | 863 ± 178 | Radiatum | 165 ± 37 | 1312 ± 288 |
| Layer V - VI | 192 ± 52 | 1394 ± 312 | CA4 |  |  |
| White Matter | 60 ± 19 | 398 ± 155 | Oriens | 169 ± 42 | 1367 ± 211 |
| Periamygdalar Cortex |  |  | Pyramidal | 118 ± 36 | 382 ± 89 |
| Layer I - II | 284 ± 60 | 915 ± 190 | Radiatum | 176 ± 44 | 1610 ± 326 |
| Layer III - IV | 287 ± 68 | 890 ± 234 | Dentate gyrus |  |  |
| Layer V - VI | 280 ± 86 | 886 ± 234 | Granular | 105 ± 43 | 251 ± 109 |
| White Matter | 124 ± 27 | 206 ± 63 | Hilus | 152 ± 51 | 808 ± 259 |
| Amygdala |  |  | Molecular | 149 ± 42 | 1314 ± 233 |
| Basal nucleus<br>(magnocellular) | 367 ± 67 | 872 ± 238 | Subiculum |  |  |
| Basal nucleus<br>(parvocellular) | 348 ± 69 | 900 ± 205 | Oriens | 179 ± 39 | 1348 ± 325 |
| Central nucleus | 380 ± 85 | 1051 ± 195 | Pyramidal | 160 ± 34 | 599 ± 106 |
| Cortical nucleus | 323 ± 50 | 842 ± 217 | Radiatum | 190 ± 39 | 1247 ± 242 |
| Lateral nucleus | 353 ± 74 | 1028 ± 242 | Lacunosum<br>moleculare | 203 ± 47 | 1818 ± 337 |
| Striatum |  |  | Cerebellum |  |  |
| Caudate | 487 ± 62 | 921 ± 200 | Granular layer | 245 ± 57 | 160 ± 79 |
| Putamen | 472 ± 62 | 751 ± 174 | Molecular layer | 259 ± 29 | 1715 ± 70 |
| Globus pallidus | 215 ± 24 | 1215 ± 162 | Whitte layer | 196 ± 36 | 48 ± 24 |
| Anterior commissure | 110 ± 51 | 350 ± 98 |  |  |  |
| Basal forebrain |  |  |  |  |  |
| Nucleus basalis<br>(Meynert) | 154 ± 56 | 1569 ± 218 |  |  |  |

Data are mean ± SEM values. n=8.

Regarding the basal ganglia, the globus pallidus showed high S1P_1_ activity (1215 ± 162 nCi/g t.e.; figure 4C), but the striatum, including the caudate and putamen nuclei, showed more moderated stimulations (921 ± 200 nCi/g t.e.; 751 ± 174 nCi/g t.e.; respectively; figure 4B). The anterior commissure, constituted by white matter fibers, was sparsely stimulated by the S1P_1_ specific agonist (350 ± 98 nCi/g t.e.; figure 4C). In addition, nbM was measured in the basal forebrain and showed high stimulation (1569 ± 218 nCi/g t.e.; figure 4C) (Table 3).

In the amygdala, we found that the activity of S1P_1_ receptor was homogenous. Despite this homogeneity, different areas could be distinguished, as central (1051 ± 195 nCi/g t.e.; figure 4D) and lateral nucleus (1028 ± 242 nCi/g t.e.; figure 4D) that were more stimulated than cortical nucleus (842 ± 217 nCi/g t.e.; figure 4D) or basal nucleus, including the magnocellular and parvocellular parts (872 ± 238 nCi/g t.e.; 900 ± 205 nCi/g t.e. respectively; figure 4D) (Table 3).

In the hippocampus, the pyramidal layer was sparsely stimulated, while the *oriens* and *radiatum* layers were strongly stimulated. Furthermore, in the dentate gyrus, the molecular layer showed a high S1P_1_ activity, but the granular layer was sparsely stimulated. About the *lacunosum moleculare*, it was strongly stimulated (1818 ± 337 nCi/g t.e.; figure 4F) (Table 3).

Finally in the cerebellum, the molecular layer showed a high activity (1715 ± 70 nCi/g t.e.; figure 4E), yet the granular layer was sparsely stimulated (160 ± 79 nCi/g t.e.; figure 4E) (Table 3).

Overall, no differences were found between S1P_1_ receptor activity when comparing male and female samples in all areas analyzed.

### 3.4 Anatomical comparison of G**_α_**_i/o_-proteins activated by the S1P_1_ receptor agonist CYM-5442 in rodents and human brain

In summary, the activity of the S1P_1_ receptor was conserved in human compared to the rat brain (Figure 5 and S3). In the cortex, the layers III - IV were strongly stimulated in both rodents (mouse and rat) and in human, but the layers V – VI showed a higher activity in both rodent brains than in human brain (Figure 5A). Regarding the striatum, in human tissue we could differentiate the caudate and putamen nucleus, in rodents, to the contrary, it is not possible, but the activity of the S1P_1_ receptor was similar in the striatal areas of humans and rodents. Moreover, the nucleus basalis magnocellularis in rodent brains and the nbM in human, which are assimilated to be comparable, both showed one of the highest measurements of S1P_1_ activity in the CNS (Figure 5B). About the hippocampus, in mouse brain the intensity of the stimulation was lower than in rat brains. Nevertheless, the S1P_1_ stimulation pattern was conserved in all the samples, being strongly stimulated in *oriens* and *radiatum* layers, more than in pyramidal layer (Figure 5C). Finally, in the cerebellum the molecular layer was the most stimulated layer in this area in both rodent and human brains, although the rat stimulation pattern is more similar to that observed in human (Figure 5D).

**Figure 5.**
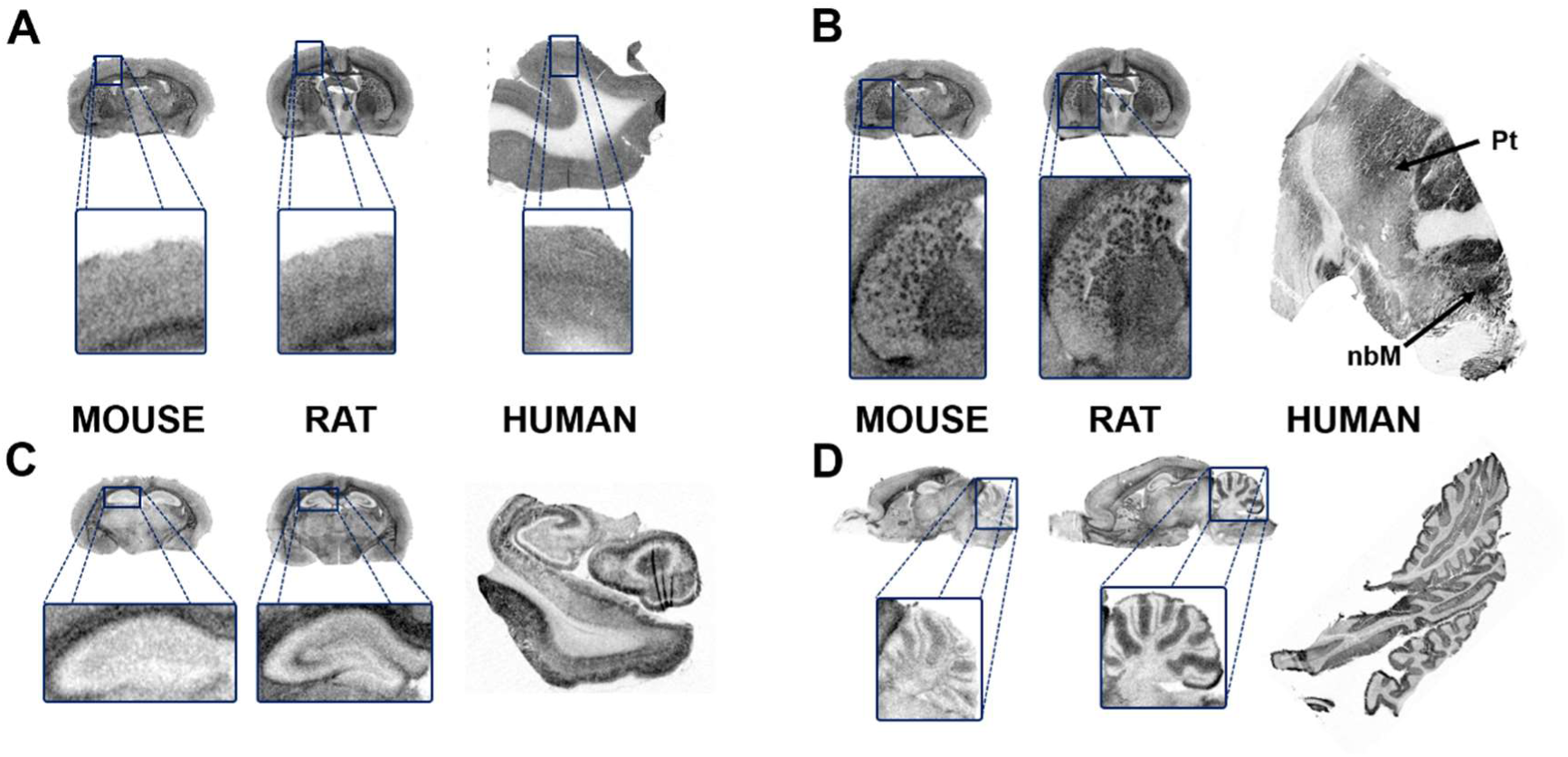
Comparative autoradiograms showing the [^35^S]GTPγS binding stimulated by the specific agonist of S1P_1_ receptor, CYM-5442, (10µM) in mouse, rat and human. Note in cortical areas that the layers V – VI show a higher activity in rodent brains than in human brain (A). Mouse, rat and human striatum and basal forebrain (Pt: putamen; nbM: nucleus basalis of Meynert) (B). Mouse, rat and human hippocampus, although the stimulation pattern is conserved in rodent brain, in mouse brain is much lower the activity of S1P_1_ receptor (C). Mouse, rat and human cerebellum, showing that the rat S1P_1_ activity is more similar to the human (D).

In general, the activity of the S1P_1_ receptors mediated by CYM-5442 showed a similar pattern in human and rat brains, including the basalocortical pathway that regulates learning and memory processes but also in striatum, hippocampus or cerebellum. The present results indicate the preference of the use of rat models to the study of S1P_1_ receptors activity in human neurodegenerative diseases, such as Alzheimer’s disease.

## 4. Discussion

The present study quantifies the anatomical distribution of the S1P_1_ receptor activity in rat, mouse and human brain to describe the similarities and differences that could further enlighten the understanding of the physiology of this type of neurolipid receptor in the mammals brain. For these purpose, we used the [^35^S]GTPγS assay to locate the distribution of the S1P_1_ receptor-mediated activity in presence of selective agonists. Firstly, we compared the activation of the S1P_1_ coupled G_αi/o_ receptors by using both S1P endogenous ligand and CYM-5442 S1P_1_ selective agonist, revealing a higher stimulation induced by CYM-5442. However, other authors have also found a good stimulation with the S1P endogenous ligand (Waeber and Chiu, 1999), or with SEW2871, another S1P_1_ receptor agonist (Sim-Selley et al., 2009).

The observed distribution of the S1P_1_ receptor activity in the brain of rat and mouse were quite similar, a wide distribution and high intensity of S1P_1_ receptor-mediated activity in the CNS was observed. High net stimulations of G_αi/o_ proteins mediated by S1P_1_ receptor were recorded at the rostral migratory stream (RMS) of the olfactory bulb (OB). In the postnatal brain, neuroblasts have to migrate long distances to the OB, for which they express S1P_1_. In the OB, down-regulation of S1P_1_ receptors facilitates radial migration of neuroblasts to the outer layers of the OB (Alfonso et al., 2015).

Regarding the cerebral cortex, the S1P_1_ activity was similar for all the analysed different cortical areas. Nevertheless, the distribution within the different layers followed a gradient from low to high from layers I - II to layers V – VI. Compared to the distribution of the activity of other receptors for neurolipids such as the CB_1_ cannabinoid receptor, the cortical activity of the S1P_1_ receptor was even higher, but its distribution was similar (Herkenham et al., 1991; Sim et al., 1996). In this regard, the S1P system is regulated in both patient cortex and animal models of Alzheimer’s disease, such as 5xFAD, suggesting the relevance of this system in cognitive degeneration (Couttas et al., 2014; Jung et al., 2023).

Although S1P_1_ receptor activity is predominant in the grey matter, it is also high in some areas of myelinated white matter, such as the corpus callosum. Indeed, S1P_1_ receptor regulates oligodendrocytes differentiation and myelination during development, and also regulates the expression of myelin basic protein in *corpus callosum* (Dukala and Soliven, 2016). Moreover, S1P_1_ agonists such as fingolimod, CYM-5442 and ponesimod, protect against toxin-induced demyelination processes in experimental models of multiple sclerosis through S1P_1_ receptor (Kihara et al., 2022; Kim et al., 2018).

The basal ganglia, including the striatum and its efferent parts, the globus pallidus and substantia nigra, also showed relevant S1P_1_ activity. The S1P_1_ receptors in these brain areas may be responsible for the effects in the regulation of motor activity in animals treated with S1P_1_ receptor agonist SEW2871 (Sim-Selley et al., 2009). Furthermore, the treatment with fingolimod in rodent models of Parkinson’s disease is able to reduce the dopaminergic neuronal damage induced by rotenone and 6-hydroxydopamine toxins (Zhao et al., 2017).

The amygdala also showed high levels of S1P_1_ activity in the present study. Therefore, this receptor could have modulating functions in anxiety-like behaviors. In this brain area the S1P is modulating the dopaminergic system as well, and seems to be involved in the induction of anxiety during stress (Jang et al., 2011).

One of the highest activities for S1P_1_ receptor was measured in the nucleus basalis magnocellularis (B), as well as in surrounding areas included in the basal forebrain. Although the physiological role of sphingolipids in basal forebrain is largely unknown, it has been proposed that the GM1 ganglioside could share the nerve grow factor (NGF) signaling pathway, being described the NGF as activator of the S1P production through the p75^NTR^ present in basal forebrain cholinergic neurons (BFCN) (Garofalo and Cuello, 1995; Zhang et al., 2006).

Regarding the hippocampus, S1P_1_ activity was similar in CA1, CA2 and CA3 regions, where the *radiatum* and *oriens* layers showed the highest S1P_1_ activities. Likewise, the molecular and polymorphic layers showed higher stimulation than granular layer in the dentate gyrus. In this regard, during developmental stages, the S1P_1_ receptor induces pleiotropic functions during neurogenesis, observing morphological changes in hippocampal-derived neural progenitor cells (Harada et al., 2004).

Concerning the cerebellum, we found a high S1P_1_ activity in the molecular layer of the cerebellar cortex, which corresponds to the area where are located the stellate and basket cells together with the dendritic arborization of the Purkinje cells. All of them are supported by the specialized radial glia known as Bergmann’s glial cells, where, the S1P hydrolyzing enzyme, LPP3, is highly expressed during the development stages (López-Juárez et al., 2011).

As mentioned before, the activity mediated by S1P_1_ receptor in *postmortem* human brain tissue was also analyzed, and it was compared with the above described for the most widely used rodent experimental models for the study of the CNS; i.e., rat and mouse. The anatomical distribution of the S1P_1_ receptor in the human brain has been poorly studied and restricted to few immunohistochemical studies to determine the cellular distribution of this receptor (Nishimura et al., 2010), or the recently published study using the new [^3^H]CS1P1 radioligand (Jiang et al., 2021). As indicated, we applied the same measurement approach used in rat and mice, to measure the S1P_1_ activity in *postmortem* human brain samples from different representative areas with interest for distinct neurodegenerative diseases, including frontal and temporal cortices, striatum, basal forebrain, amygdala, hippocampus and cerebellum.

The S1P_1_ activity patterns obtained in the human cortical areas were similar to those previously observed in rodent brains. However, there were few peculiarities in terms of stimulation in the different cortical areas. The layers III – VI of the frontal and temporal cortices showed higher stimulation than layers I – II and V – VI, while the layers V – VI of the entorhinal cortex were highly stimulated compared to layers I – II and III – IV. This change of pattern compared to that found in rodents may be related to the different distribution of cells in human cortical areas (Ribeiro et al., 2013; Zilles et al., 2004). In general, the cortical S1P_1_ activity was higher in the deeper layers of rodents compared to human brain. In addition, the white matter seems to be also more stimulated in rodents than in humans, possibly due to the different packing of the white matter described in rodents as opposed to the greater amplitude found in humans thanks to the gyrification of the cortex (Mota et al., 2019; Ventura-Antunes et al., 2013).

In relation to the activity of S1P_1_ receptor in the basal ganglia, it was high at globus pallidus while the striatum, including the caudate and putamen nuclei, showed moderate levels, the latter, possibly due to the contribution of the highly myelinated striatopallidal fibers. Moreover, the nbM located at the basal forebrain, showed a high stimulation in a similar way to that found for the BFCN in rat. Interestingly, both structures, the nucleus basalis magnocellularis (B) in rodent brains and the nbM in human, showed the highest measurements of S1P_1_ activity, probably indicating that S1P_1_ receptors are regulating their physiological functions. Moreover, in view of these results it can be inferred that the rodents are a good model to study the role of this subtype of S1P receptor in this area, very probably modulating the cholinergic signaling that governs the memory and learning controlled by the basalocortical pathway. In the same way, the activity of S1P_1_ receptor was similar in the striatal areas of rodents and humans, therefore the rodents could constitute also a good model to study the impairment of the nigro-striatal pathway in motor-related diseases including Parkinson’s and Hungtinton’s disease.

In the hippocampus, the S1P_1_ activity pattern resembles that seen in the rat but not in the mouse brain. In this sense, the *oriens* and *radiatum* layers showed high levels, while in the pyramidal layer was more weak. In addition, the molecular layer of the dentate gyrus showed a high S1P_1_ activity, but in the granular layer was low. It is assumed that the neurogenesis in humans, as well as in rodents brain, also occurs in this region. This process is regulated by glutamatergic inputs, which as above mentioned, can be regulated by S1P_1_ receptor activity in this area (Kempermann et al., 2015).

In summary, the described anatomical distributions of S1P_1_ receptor mediated-activity in rat, mice and human by [^35^S]GTPγS assay, are similar to those obtained by immunohistochemical (Nishimura et al., 2010), or radioligand binding assays (Jiang et al., 2021). In general, the activity of the S1P_1_ receptors elicited by CYM-5442 showed a more similar pattern in human and rat brains than in mouse. Since the S1P_1_ activity is more similar in rat and human brain at pathways involved in control of learning and memory, motor and nociception, it seems more appropriate the use of rats as animal models to study the S1P_1_ receptor in neuropathologies affecting those functions, i.e., Alzheimer’s, Parkinson’s, Huntington’s diseases or multiple sclerosis.

## 5. Conclusion

S1P_1_ receptor is one of the most abundant and efficient GPCRs coupled to G_αi/o_ proteins in the human and rodent brain, with the highest activity in basal forebrain, which indicates an important role in learning and memory processes. Moreover, it could be signaling a vast amount of different biological processes by regulating or modulating other systems, resulting in very relevant roles in the control of brain functioning. Furthermore, the rat would be a more preferable experimental model than mice to extrapolate the S1P_1_ receptor-mediated responses to humans.

## Acknowledgements

Technical and human support provided by University of the Basque Country (UPV/EHU), Ministry of Economy and Competitiveness (MINECO), Basque Government (GV/EJ), European Regional Development Fund (ERDF), and European Social Fund (ESF) is gratefully acknowledged. J.M.-G. is the recipient of Margarita Salas fellowship funded by the European Union-Next Generation EU. G.P.-C is the recipient of predoctoral fellowship funded by the University of the Basque Country (UPV/EHU).

## ACRONYMS AND ABBREVIATIONS

B: Nucleus basalis magnocellularis
BFCN: Basal forebrain cholinergic neurons
BSA: Bovine serum albumin
CNS: Central nervous system
Cx: Cortex
DTT: DL-dithiothreitol
EDG: Endothelial differentiation gene
GDP: Guanosine 5’-diphosphate
GPCR: G protein-coupled receptor
GTPγS: Guanosine 5’-O-(3-thiotriphosphate)
LPA: Lysophosphatidic acid
mcp: middle cerebellar peduncles
nbM: Nucleus basalis of Meynert
OB: Olfactory bulb
p75^NTR^: p75 pan-neurotrophin receptor
Pt: Putamen
RMS: Rostral migratory stream
S1P: Sphingosine 1-phosphate
SEM: Standard error of the mean
Str: Striatum
Th: Thalamus

