## Supplemental Material for "Localization of S1P_1_ Receptor Signaling in the rat, mouse and human Central Nervous System"

### SUPPLEMENTARY MATERIAL

#### S1. Preliminary studies of $S1P_1$ receptor-mediated activity by [$^{35}$S]GTPγS autoradiography assay in rodent CNS

It was noted that, despite the specificity of the $S1P_1$ agonist, the CYM-5442 was able to stimulate the binding of [$^{35}$S]GTPγS more than S1P in all areas of the CNS that were analyzed. About the gray matter areas, the molecular layer of the cerebellum was the area in which the net stimulation evoked by CYM-5442 over the basal levels was higher (2644 ± 275 nCi/g t.e.) in comparison with the stimulation by S1P (109 ± 74 nCi/g t.e.). This superior stimulation by CYM-5442 was also observed in other measured areas of gray matter, such as olfactory bulb (CYM-5442: 636 ± 107 nCi/g t.e.; S1P: 30 ± 84 nCi/g t.e.), cortex (CYM-5442: 1251 ± 95 nCi/g t.e.; S1P: 59 ± 72 nCi/g t.e.), striatum (CYM-5442: 775 ± 120 nCi/g t.e.; S1P: 99 ± 39 nCi/g t.e.), globus pallidus (CYM-5442: 431 ± 100 nCi/g t.e.; S1P: -110 ± 85 nCi/g t.e.), thalamus (CYM-5442: 481 ± 97 nCi/g t.e.; S1P: 78 ± 57 nCi/g t.e.), hippocampus (CYM-5442: 684 ± 86 nCi/g t.e.; S1P: 8. ± 49 nCi/g t.e.), inferior colliculus (CYM-5442: 425 ± 143 nCi/g t.e.; S1P: 36 ± 43 nCi/g t.e.), superior colliculus (CYM-5442: 415 ± 115 nCi/g t.e.; S1P: -29 ± 56 nCi/g t.e.) and granular layer of the cerebellum (CYM-5442: 348 ± 39 nCi/g t.e.; S1P: 109 ± 74 nCi/g t.e.). Furthermore, the same effect was also observed in some areas of white matter that were analyzed such as corpus callosum (CYM-5442: 236 ± 99 nCi/g t.e.; S1P: 87 ± 58 nCi/g t.e.), fimbria of the hippocampus (CYM-5442: 45 ± 54 nCi/g t.e.; S1P: -54 ± 31 nCi/g t.e.), cerebellar white matter (CYM-5442: 21 ± 36 nCi/g t.e.; S1P: -30 ± 39 nCi/g t.e.) and brainstem (CYM-5442: 49 ± 47 nCi/g t.e.; S1P: -36 ± 67 nCi/g t.e.) (Figure S1).

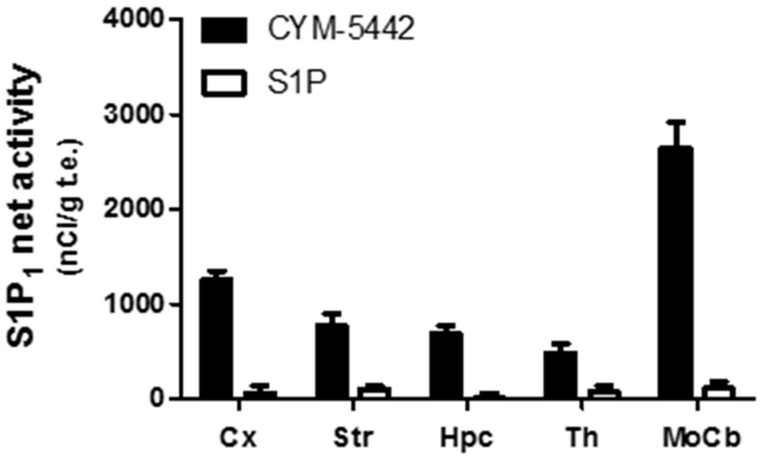


Figure S1. [$^{35}$S]GTPγS binding stimulated by CYM-5442 (10 µM) and S1P (10 µM) in different brain areas expressed as nCi/g tissue equivalent (t.e.). Note that the specific S1P$_1$ agonist (CYM-5442) evoked higher activity than the endogenous ligand, the S1P. Abbreviations: Cx: Cortex; Str: Striatum; Hpc: Hippocampus; Th: Thalamus; MoCb: Molecular layer of the cerebellum. Data are mean ± S.E.M. n=8.

The compound W146, which is a selective antagonist of S1P$_1$ receptor, was used to determine the S1P$_1$ receptor-mediated specificity of the stimulations when were co-incubated in the presence of agonists. It was noteworthy that W146 blocked the stimulations in all areas for both compounds, CYM-5442 or S1P (Figure S2E and S2F).

Moreover, we studied the possibility that the S1P$_1$ receptor activity was contributing in some way to the basal binding inherently detected by [$^{35}$S]GTPγS binding technique. If S1P$_1$ receptor was contributing to the basal activity in the assayed conditions, W146 would also block the stimulation of the basal binding, assuming that W146 is specific of S1P$_1$ receptors at the concentration used. With this in mind, we incubated a consecutive slice with W146 in absence of any of the agonists (Figure S2D); when the result was compared to basal binding (Figure S2A), and no significant differences were found, discarding this possibility.

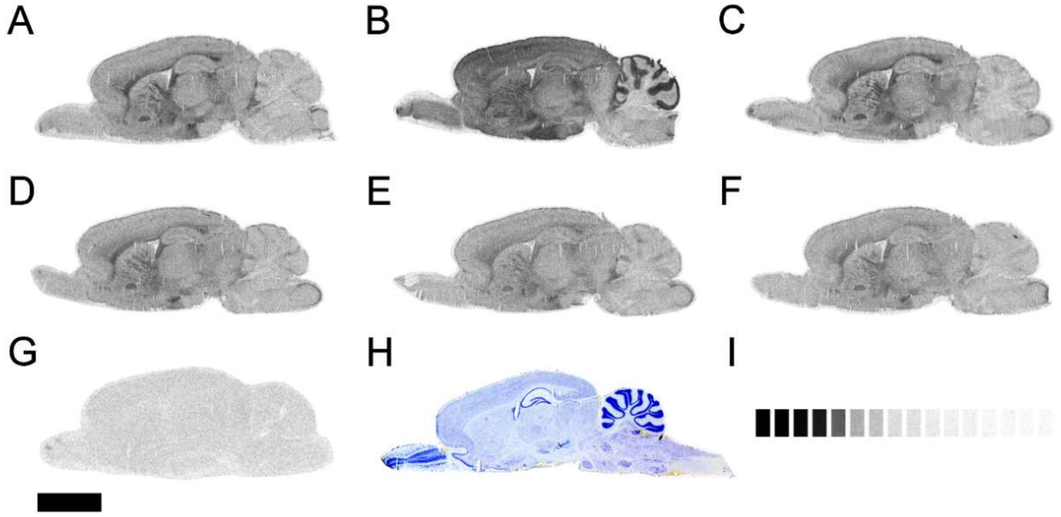

Figure S2. Representative autoradiograms corresponding to consecutive sagittal sections from control rats showing [$^{35}$S]GTPγS basal binding (A), stimulated by the specific $S1P_1$ agonist CYM-5442 (10 µM) (B), stimulated by the agonist S1P (10 µM) (C), basal binding in presence of the specific antagonist of $S1P_1$, W146 (10 µM) (D), CYM-5442 (10 µM) stimulation antagonized with W146 (10 µM) (E), S1P (10 µM) stimulation antagonized with W146 (10 µM) (F), the non-specific binding was defined in the presence of GTPγS (10 µM) (G). Thionine staining (H). [$^{14}$C]-Standard (35000 – 0 nCi/g t.e.) (I). Scale bar = 6 mm.

In summary, taking all of these results into account, CYM-5442 elicited higher $S1P_1$ receptor-mediated stimulations than S1P in the rat CNS. Consequently, we hereafter used CYM-5442 as the agonist of choice to analyze $S1P_1$ receptor-mediated activity using the [$^{35}$S]GTPγS binding assay, and W146 as antagonist.

**S2. Localization of S1P1 receptor-mediated activity by [$^{35}$S]GTPγS autoradiography assay in the mouse CNS**

We also carried out the mapping of the $S1P_1$ receptor-mediated activity by [$^{35}$S]GTPγS autoradiography assay in mouse brain. In this sense, we collected slices of the brain cutting them rostro-caudally every 500 µm to obtain an exhaustive mapping of the $S1P_1$ receptor stimulation in the different structures present in the rat brain. The obtained stimulations of [$^{35}$S]GTPγS binding by CYM-5442 (10 µM) showed that the activity of $S1P_1$ receptor was abundant along the brain in a similar way to that found for the rat brain (Table S2).

**Table S2**. Net stimulation of [$^{35}$S]GTPγS binding by the agonist of $S1P_1$ receptor, CYM-5442 (10 μM), in mouse brain (nCi/g t.e).

| AREA | $S1P_1$ net stimulation (nCi/g t.e.) |
|---|---|
| **CEREBRUM** | |
| Olfactory bulb | |
| Rostral migratory stream | 1219 ± 166 |
| External plexiform layer | 448 ± 52 |
| Internal plexiform layer | 951 ± 107 |
| Glomerular layer | 370 ± 69 |
| Lateral olfactory tract | 1779 ± 137 |
| Cerebral Cortex | |
| Frontal | 1439 ± 128 |
| Cingulate | 1381 ± 91 |
| Motor | 1495 ± 75 |
| Somatosensory | 1617 ± 122 |
| Piriform | 941 ± 89 |
| Layer I - II | 932 ± 95 |
| Layer III - IV | 953 ± 69 |
| Layer V - VI | 1059 ± 105 |
| Agranular insular | 1385 ± 104 |
| Granular insular | 1527 ± 205 |
| Intermediate endopiriform nucleus | 1073 ± 125 |
| Dorsal endopiriform nucleus | 1027 ± 117 |
| Claustrum | 1833 ± 128 |
| Corpus callosum | 3072 ± 180 |
| Forceps minor of corpus callosum | 1320 ± 174 |
| Basal ganglia | |
| Striatum | 1534 ± 126 |
| Striatum fibers | 2435 ± 200 |
| Internal capsule | 2480 ± 303 |
| Anterior commissure, anter part | 2729 ± 162 |
| Accumbens nucleus | |
| Core | 1149 ± 88 |
| Shell | 1038 ± 160 |
| Globus pallidus | 2623 ± 323 |
| Ventral pallidum | 1394 ± 253 |
| Substantia nigra | 1598 ± 178 |
| Amygdala | |
| Basolateral amygdaloid nucleus | 1733 ± 528 |
| Central amygdaloid nucleus | 1880 ± 408 |
| **DIENCEPHALON** | |
| Basal forebrain | |
| Basal nucleus magnocellularis | 2920 ± 326 |
| Medial septal nucleus | 1561 ± 244 |
| Hippocampus | |
| CA1 | 929 ± 129 |
| Oriens layer | 1111 ± 120 |
| Pyramidal layer | 527 ± 39 |
| Radiatum layer | 899 ± 62 |
| CA2 | 622 ± 30 |
| Oriens layer | 883 ± 65 |
| Pyramidal layer | 339 ± 47 |
| Radiatum layer | 770 ± 73 |
| CA3 | 763 ± 90 |
| Oriens layer | 722 ± 51 |
| Pyramidal layer | 571 ± 35 |
| Radiatum layer | 733 ± 77 |
| Dentate gyrus | 590 ± 56 |
| Polymorph layer | 562 ± 92 |
| Granule cell layer | 375 ± 52 |
| Molecular layer | 723 ± 98 |
| Lacunosum moleculare layer | 1101 ± 102 |
| Stratum lucidum | 1147 ± 84 |
| **BRAINSTEM** | |
| Superior Colliculus | |
| Superficial gray layer | 895 ± 170 |
| Optic nerve layer | 2226 ± 171 |
| Intermediate gray | 1734 ± 225 |
| Intermedial white layer | 1553 ± 187 |
| Deep gray layer | 1781 ± 126 |
| Deep white layer | 1375 ± 216 |
| Commisure | 1839 ± 226 |
| External cortex of inferior colliculus | 2362 ± 259 |
| Perioacueductal gray | 1473 ± 196 |
| Median raphe nucleus | 1342 ± 183 |
| Pontine nucleus | 919 ± 129 |
| Locus coeruleus | 1460 ± 209 |
| **CEREBELLUM** | |
| Cerebellar cortex | |
| Molecular layer | 595 ± 46 |
| Granular layer | 356 ± 67 |
| White matter | 1024 ± 155 |

Data are mean ± SEM values. n=8.

**S3. Comparison among basal, stimulated, antagonised and non-specific binding in rat, mouse and human slices.**

The compound W146, which is a selective antagonist of $S1P_1$ receptor, blocked the stimulations mediated by CYM-5442 in all tissues analyzed (Figure S3).

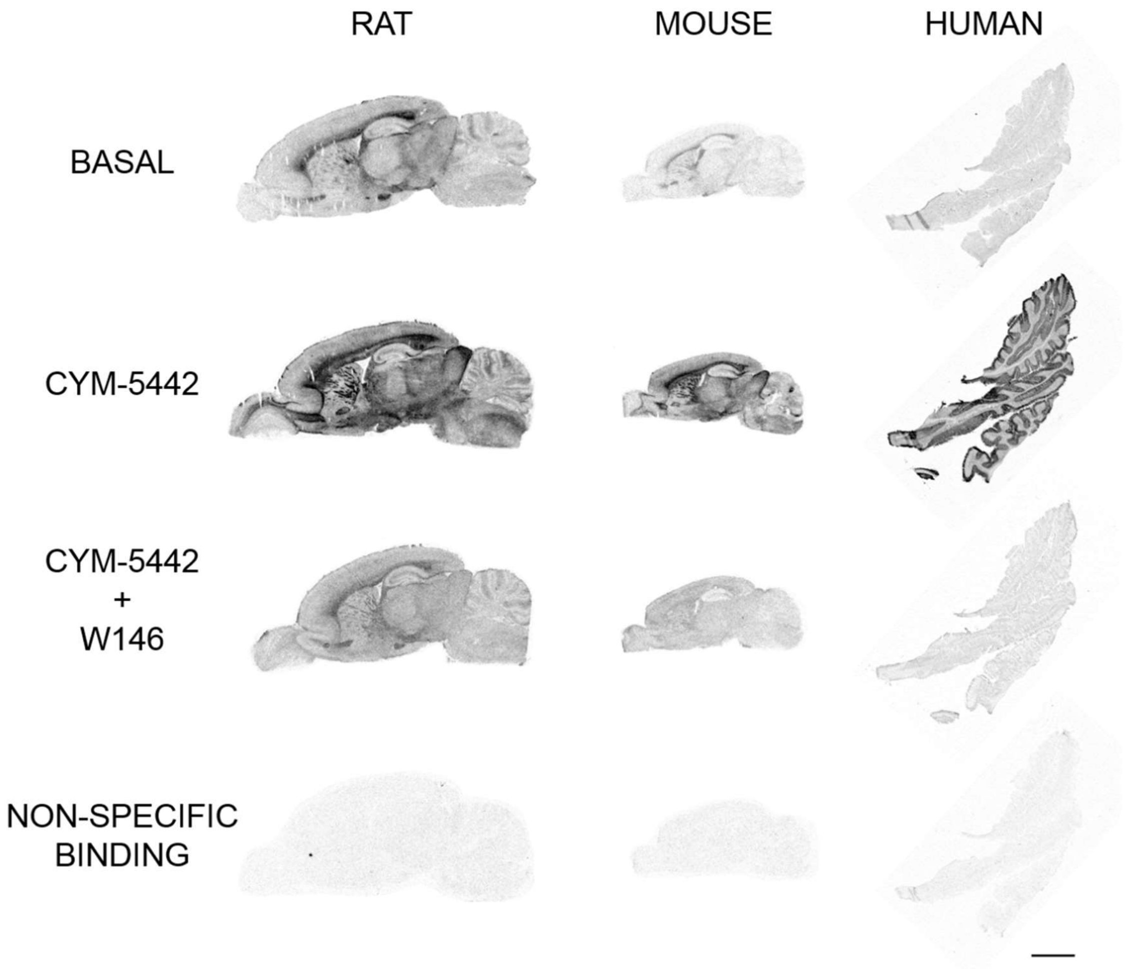


Figure S3. Representative autoradiograms corresponding to consecutive sagittal sections from control rats, mouse and human cerebellum showing [$^{35}$S]GTPγS basal binding, stimulated by the specific $S1P_1$ agonist CYM-5442 (10 μM), CYM-5442 (10 μM) stimulation antagonized with W146 (10 μM) and the non-specific binding was defined in the presence of GTPγS (10 μM). Scale bar = 6 mm.
